# DiffusionST: A deep generative diffusion model-based framework for enhancing spatial transcriptomics data quality and identifying spatial domains

**DOI:** 10.1101/2025.06.12.659243

**Authors:** Yaxuan Cui, Yang Cui, Ruheng Wang, Zheyong Zhu, Xin Zeng, Kenta Nakai, Feifei Cui, Zilong Zhang, Hua Shi, Yan Chen, Xiucai Ye, Tetsuya Sakurai, Leyi Wei

## Abstract

Recent advancements in spatial transcriptomics technology have generated substantial volumes of spatial transcriptome data. However, the quality of this data is often compromised due to the limitations of current sequencing technologies. To address this issue, DiffusionST proposes a method for imputing spatial transcriptomics data and clustering the imputed data. The method employs a graph convolutional network (GCN) model combined with a newly designed loss function, denoising data using the zero-inflated negative binomial (ZINB) distribution, and data enhancement through a diffusion model to improve clustering accuracy. DiffusionST demonstrates superior clustering accuracy compared to six of the most popular spatial transcriptomics clustering algorithms. DiffusionST also excels in data imputation when compared to five single-cell RNA sequencing (scRNA-seq) imputation algorithms. Additionally, DiffusionST’s robustness against noise is quantitatively validated by manually introducing random dropout noise into the dataset, where our model significantly enhances the quality of spatial transcriptomic data. Moreover, DiffusionST is well-suited for high-resolution spatial transcriptomics data and has been demonstrated, through survival analysis and cell-cell communication studies, to dissect spatial domains within breast cancer tissues. These findings provide strong evidence of DiffusionST’s efficacy in handling spatial transcriptomic data especially with strong noise, making it a valuable tool in this field.

## Introduction

Spatial transcriptomics has garnered significant attention from researchers as it enables gene expression analysis across numerous spatial locations within a tissue[1]. Compared to traditional whole-transcriptome sequencing, spatial transcriptomics provides more detailed information about tissue structure, helping researchers better understand the spatial distribution characteristics of gene expression in organisms and their roles in physiological and pathological processes[2]. Spatial transcriptomics provides researchers with tissue image data, spatial location data, and corresponding gene expression data. The field of spatial transcriptomics encompasses a collection of emerging techniques developed in recent years[3]. These techniques include those based on single molecule fluorescent in situ hybridization, such as seqFISH[4], seqFISH+[5], and MERFISH[6]; as well as those based on in situ mRNA capturing followed by high-throughput sequencing techniques, such as Stereo-seq[7], Slide-seq[8], and 10x Visium[9].

Recently proposed spatial transcriptomics clustering methods consider the similarity between neighboring spots, thereby better explaining the spatial dependency of gene expression. For example, BayesSpace[10] uses a probability density method for clustering, introducing priors to assign higher weights to physically proximate spots. SpaGCN[11] uses an undirected weighted graph to represent the dependencies of spatial data, where spatial location, histological images, and gene expression are involved in the graph construction. STAGATE[12] uses a graph attention autoencoder framework, integrating spatial information and gene expression profiles to learn low-dimensional representations of spatial transcriptomics data spots. DeepST[13] employs denoising autoencoders and GNN autoencoders to jointly infer enhanced latent embeddings of spatial transcriptomics data. GraphST[14] uses a graph self-supervised contrastive learning framework to learn latent embeddings of spatial transcriptomics data. SEDR[15] is a deep masked autoencoder network used for learning latent representations, while the other is a variational graph autoencoder network for embedding spatial information. These two components are jointly optimized to generate latent representations suitable for spatial transcriptomics data analysis.

A major issue in spatial transcriptomics is the abundance of zero values in gene expression data, leading to data sparsity[16]. These zero values are mostly caused by technical defects or artificial factors, such as mRNA molecule scarcity, low capture rates, and insufficient sequencing depth[17]. Current imputation algorithms primarily utilize scRNA-seq data[18]. For example, kNN-smoothing[19] uses filtered scRNA-seq data to impute by considering the K nearest cells for each cell. SAVER[20] is a method that optimizes the entire gene expression count by using information between genes and cells to imputing zero values and improve the data quality of all genes. MAGIC[21] applies Markov affinity-based graph techniques to share information between similar cells, optimizing the gene expression matrix and imputing zero values. The deep count autoencoder network[22] (DCA) captures nonlinear gene-gene correlations by introducing a zero-inflated negative binomial noise model while considering the count distribution, over-dispersion, and sparsity of the data to infer missing values. ALRA[23] can selectively impute missing values through non-negativity and correlation structure, effectively maintaining biologically meaningful zero values while addressing “dropout events”.

Previously proposed imputation algorithms were designed for scRNA-seq data and did not consider the specific characteristics of spatial transcriptome data, such as spatial positional information. Therefore, these algorithms are not suitable for use in spatial transcriptome data. There is an urgent need to design imputation algorithms adapted to spatial transcriptome data. In this study, we proposed DiffusionST, a denoising diffusion framework[25] for data enhancement and clustering of spatial transcriptomics. Inspired by image restoration, we input the spatial transcriptome’s positional information and gene expression matrix into Graph Convolutional Networks[29] (GCN) optimized with the Zero-Inflated Negative Binomial[22] (ZINB) loss function to obtain denoised data. This denoised data, which has improved quality, is then fed into a diffusion model for further imputation. These steps collectively enhance the data, facilitating more robust downstream analysis. We systematically evaluated the performance of DiffusionST. Firstly, we evaluated the accuracy of data clustering after imputation and compared it with the clustering accuracy of five popular clustering algorithms (BayesSpace, SpaGCN, STAGATE, DeepST, GraphST, SEDR). Additionally, we performed clustering accuracy comparisons by randomly initializing and adding “dropout noise” to the data. Furthermore, we compared DiffusionST-processed data with the original data and quantitively verified that our imputed data has higher data quality. Furthermore, applying DiffusionST to breast cancer spatial transcriptomic data revealed detailed spatial domains within tumor tissues, highlighting significant differences in tumor progression and immune response, and provided deeper insights into tumor heterogeneity, which can enhance cancer treatment strategies.

## Results

The diffusion model network was first proposed in 2015 and has since attracted the interest of computer vision researchers, demonstrating outstanding performance in the field of image restoration. Inspired by its algorithmic approach and the recent challenges in ST data, we propose DiffusionST. It utilizes the image restoration model to enhance spatial transcriptomic data. DiffusionST can enhance data quality from each spot containing dropout noise by learning its data distribution. DiffusionST mainly consists of two steps: Firstly, we use ‘seurat_v3’ in scanpy[30] to filter genes in the data. Then, we input the identified highly variable genes into a GCN autoencoder model with ZINB. Through this model, with the assistance of spatial information and the ZINB gene distribution, the impact of dropout noise in the data is reduced. Secondly, to further improve data quality, we reshape the expression profiles of each processed spot into images and input them into the diffusion model. The diffusion model can generate pseudo-images for ST gene expression profiles by learning the distribution process of gene expression profiles, utilizing a U-Net[31,32] with a self-attention module to enhance the network’s global modeling capability. We calculate the similarity between the pseudo-images generated by the diffusion model for each spot and the original images. Then, we select the two pseudo-images closest to the processed spot and average them to replace the processed spot. The specific reasons for selecting two pseudo-images are shown in **Table S2**. Finally, clustering is performed on the data enhanced after data augmentation (as shown in **Figure 1**).

**Figure 1.**
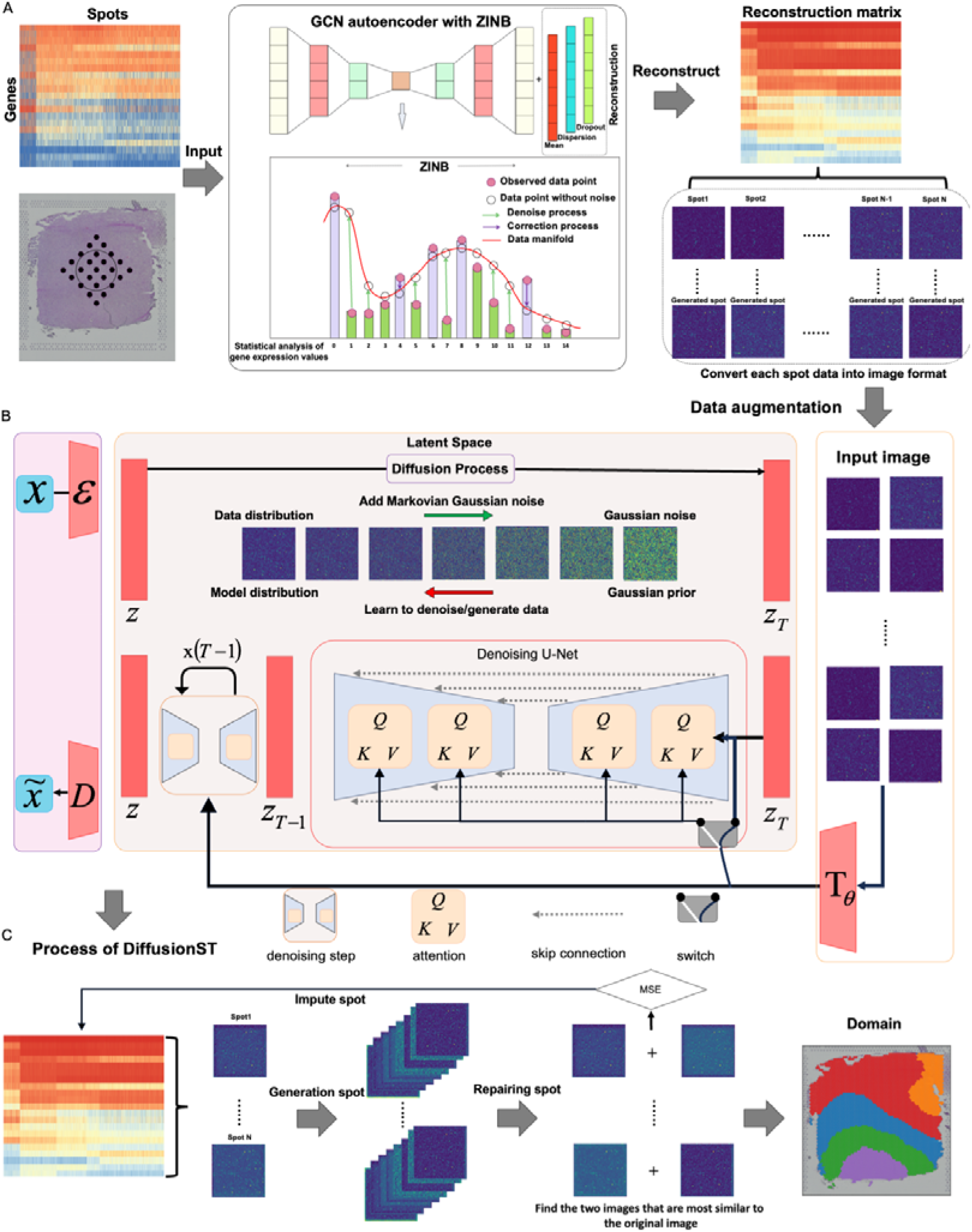
Workflow of DiffusionST. **(A)** The model learns the distribution process of gene expression profiles to generate pseudo-images for ST gene expression profiles (The generated grayscale image is visualized using the colormap tool, which is more convenient for observation compared to black-and-white images). **(B)** The similarity between the pseudo-images generated by the Diffusion model for each spot and the reconstructed spot is then calculated. The average of the two pseudo-images closest to the reconstructed spot is selected to replace the reconstructed spot. **(C)** Process of spatial transcriptomic data enhancement by the Diffusion model.

### Comparison of clustering accuracy between DiffusionST results and other popular spatial transcriptomic clustering algorithms

We compared the clustering results of the DiffusionST model after data augmentation with six of the most prominent algorithms on twelve benchmark datasets. We ran these tools using default parameter settings (see **Supplementary Note S3**). To comprehensively analyze the clustering results from a computational perspective, we compared the clustering results of twelve slice datasets using 6 evaluation metrics. Comparing Adjusted Rand Index[33] (ARI) scores on twelve slice datasets (as shown in **Figure 2A**), we found that DiffusionST ranked first in ARI scores on 6 out of 12 datasets. GraphST ranked first on 3 datasets, SEDR ranked first on 1 dataset, while STAGATE ranked first on 2 datasets. The ARI scores of BayesSpace ranged from 0.30 to 0.55 on the twelve datasets. SpaGCN’s ARI scores ranged from 0.30 to 0.57 on the 12 datasets. STAGATE’s ARI scores ranged from 0.33 to 0.60 on the 12 datasets. DeepST’s ARI scores ranged from 0.32 to 0.57 on the twelve datasets. GraphST’s ARI scores ranged from 0.43 to 0.63 on the twelve datasets. SEDR’s ARI scores ranged from 0.38 to 0.60 on the twelve datasets. DiffusionST’s ARI scores ranged from 0.43 to 0.65 on the twelve datasets. Compared to other algorithms, DiffusionST and GraphST had higher minimum ARI scores. DiffusionST demonstrated higher accuracy than GraphST on the twelve datasets (see **Figure 2B, Figure 2C and Supplementary Figure S1**). We also used other evaluation metrics to assess the accuracy of clustering (**Supplementary Note S3**).

**Figure 2.**
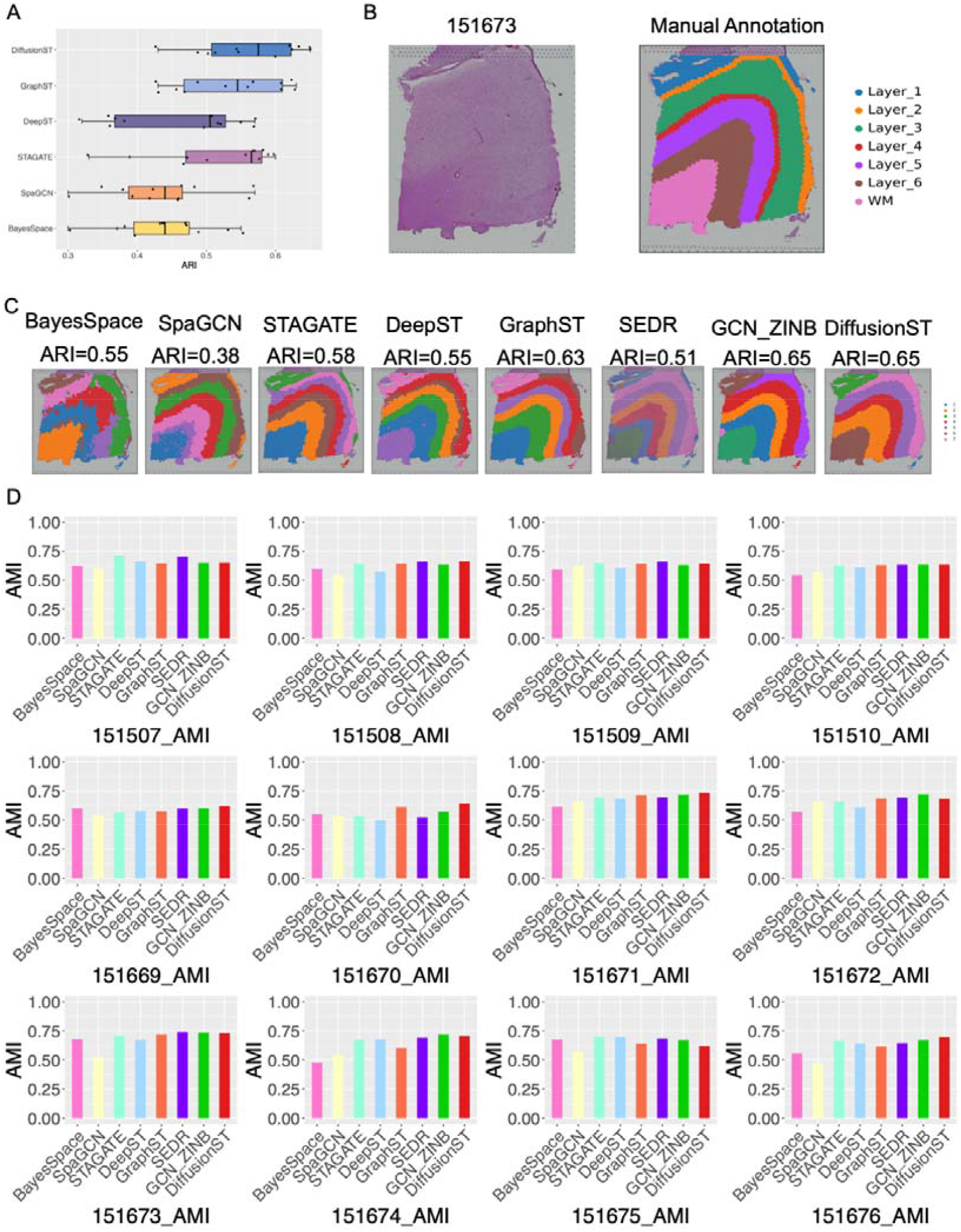
DiffusionST clustering enhances the identification of tissue structures in the dorsolateral prefrontal cortex (DLPFC). **(A)** Boxplots depicting the ARI scores of the seven methods applied to the 12 DLPFC slices. **(B)**. Histology and manual annotation images from the original study. **(C)** The clustering outcomes generated by various spatial transcriptomics techniques, encompassing BayesSpace, SpaGCN, STAGATE, DeepST, GraphST, SEDR, and DiffusionST, are presented for the 151673 slice of DLPFC dataset. Manual annotations and clustering outcomes for the remaining DLPFC slices are depicted in **Supplementary Figure S1. (D)** Comparisons of AMI scores for DiffusionST and the other six spatial transcriptomics clustering algorithms.

### DiffusionST demonstrates resilience against random initialization and dropout noise

Many machine learning and deep learning models are highly sensitive to the initialization of model parameters, and using different initializations may lead to different outputs. To better evaluate the stability of models, we conducted 10 times of clustering analyses with seven methods (DiffusionST, SEDR, GraphST, STAGATE, DeepST, SpaGCN, and BayesSpace) on twelve datasets with random initialization. We found that DiffusionST exhibits small variations in ARI scores (SD<0.06) on 12 datasets with 10 times of random initialization, demonstrating high stability. In contrast, the other five algorithmic tools show larger variations in ARI scores on 12 datasets with 10 times random initialization. (BayesSpace with SD<0.16, SpaGCN with SD<0.06, DeepST<0.07, STAGATE<0.08, GraphST<0.09, SEDR<0.07). We found that DiffusionST ranks first in ARI median scores on seven datasets. STAGATE ranks first on four datasets, and GraphST ranks first on one dataset among twelve datasets with 10 times of random initialization. We found that DiffusionST ranks first in ARI mean scores on seven datasets, STAGATE ranks first on four datasets, and GraphST ranks first on one dataset among twelve datasets with 10 times of random initialization (see **Figure 3A**).

**Figure 3.**
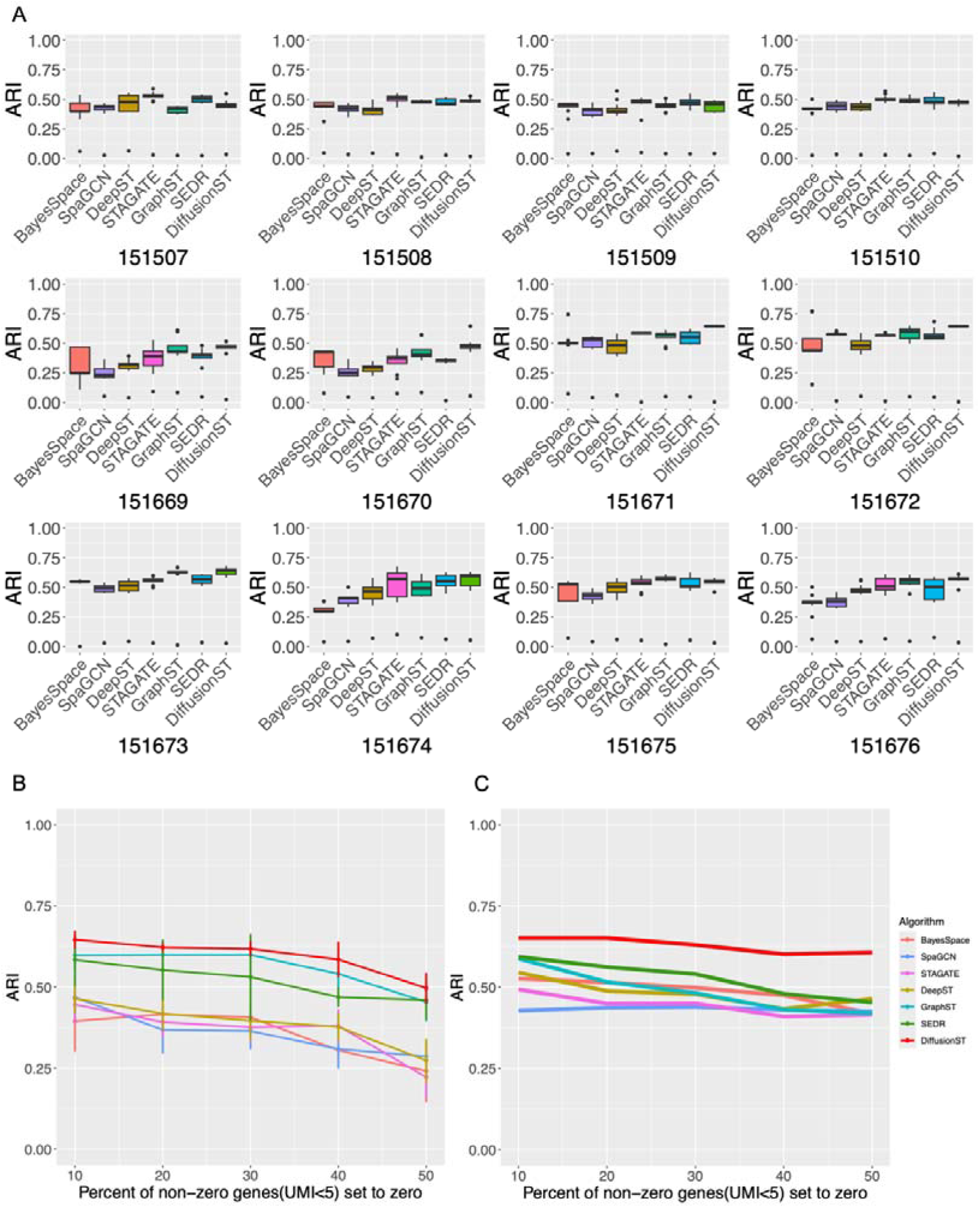
The stability assessment between DiffusionST and alternative methodologies, and robustness against dropout noise. **(A)** The ARI fluctuations arising from random initialization are examined for DiffusionST, SEDR, GraphST, STAGATE, DeepST, SpaGCN, and BayesSpace. **(B)** The performance of the seven methods is compared under varying degrees of artificial dropout noise applied to the 151673 slice (Median ARI score in 10 randomly generated replicates). **(C)** The performance of the seven methods is compared under varying degrees of artificial dropout noise applied to the 151673 slice (Mean ARI score in 10 randomly generated replicates).

Additionally, a significant issue with spatial transcriptomic data based on next-generation sequencing is the generation of “dropout noise” during the sequencing process. To further demonstrate the robustness of DiffusionST against dropout noise in spatial transcriptomics, we artificially introduced dropout noise at different proportions in the 151673 slice. Considering that most spatial transcriptomic algorithms exhibit high clustering accuracy on the 151673 slice, and dropout noise is more likely to be generated by genes with low expression levels, we introduced dropout noise by setting different percentages of non-zero elements (UMI counts <= 5) to zero. In **Figures 3B and 3C**, we repeated the process 10 times with randomly added dropout noise and calculated their clustering results using different tools to obtain the average ARI score, median ARI score, and variance. We found that DiffusionST has the highest clustering accuracy and stability. The variation in ARI scores of DiffusionST with randomly added dropout noise is relatively small (SD<0.06 for DiffusionST, SD<0.07 for DeepST, STAGATE, and GraphST, SD<0.08 for SpaGCN, SD<0.1 for BayesSpace, SD<0.13 for SEDR). Even when the dropout noise level was increased to 50%, the average ARI value of DiffusionST still reached 60, while the average ARI values of the other five tools sharply declined.

### DiffusionST can effectively enhance the quality of spatial transcriptomic data

Spatial transcriptomic data poses a significant challenge, as technical noise generated during the sequencing process leads to many dropout events in the data, severely affecting downstream analysis. DiffusionST can effectively impute and enhance the original gene expression matrix of spatial transcriptomic data using spatial information, mitigating these challenges. In the 151673 slice, we selected feature genes from the raw data for Uniform Manifold Approximation and Projection (UMAP) dimensionality reduction to two dimensions, labeled using ground truth, and then applied the same process to the data after imputation and enhancement. We observed that the scatter plot generated by UMAP of data after embedding via the DiffusionST model better separates different spot clusters compared to the original matrix (as shown in **Figure 4A**). Since we have the true labels of the data, we can indirectly demonstrate the higher data quality of the imputed and enhanced data by comparing classifications using the original expression matrix and the embedded expression matrix, thereby obtaining better downstream analysis results. In the confusion matrix before and after the imputation of the 151673 slice. In **Figure 4B**, we observed that the values on the diagonal of the matrix after embedding are higher, indicating better classification performance of the model in that category. We also noted that the values of the diagonal of the matrix after embedding are smaller, indicating better classification performance of the model in that category. In **Figure 4C**, the Precision-Recall (PR) curve of the 151673 slice, we observed that after embedding, the enhanced data compared to the original matrix, when calculating the area under the PR curve for 7 categories, the Area Under Curve Precision-Recall (AUC-PR) closer to 1, demonstrating that the data quality is better after DiffusionST processing. In **Figure 4D**, the Receiver Operating Characteristic (ROC) curve of the 151673 slice, we observed that after embedding, the enhanced data compared to the original matrix, when calculating the area under the ROC curve for 7 categories, the ROC closer to 1, demonstrating that the data quality is better after DiffusionST processing. We also conducted analysis and discussion on the 151672 spatial transcriptomics data slice (**Supplementary Note S4**).

**Figure 4.**
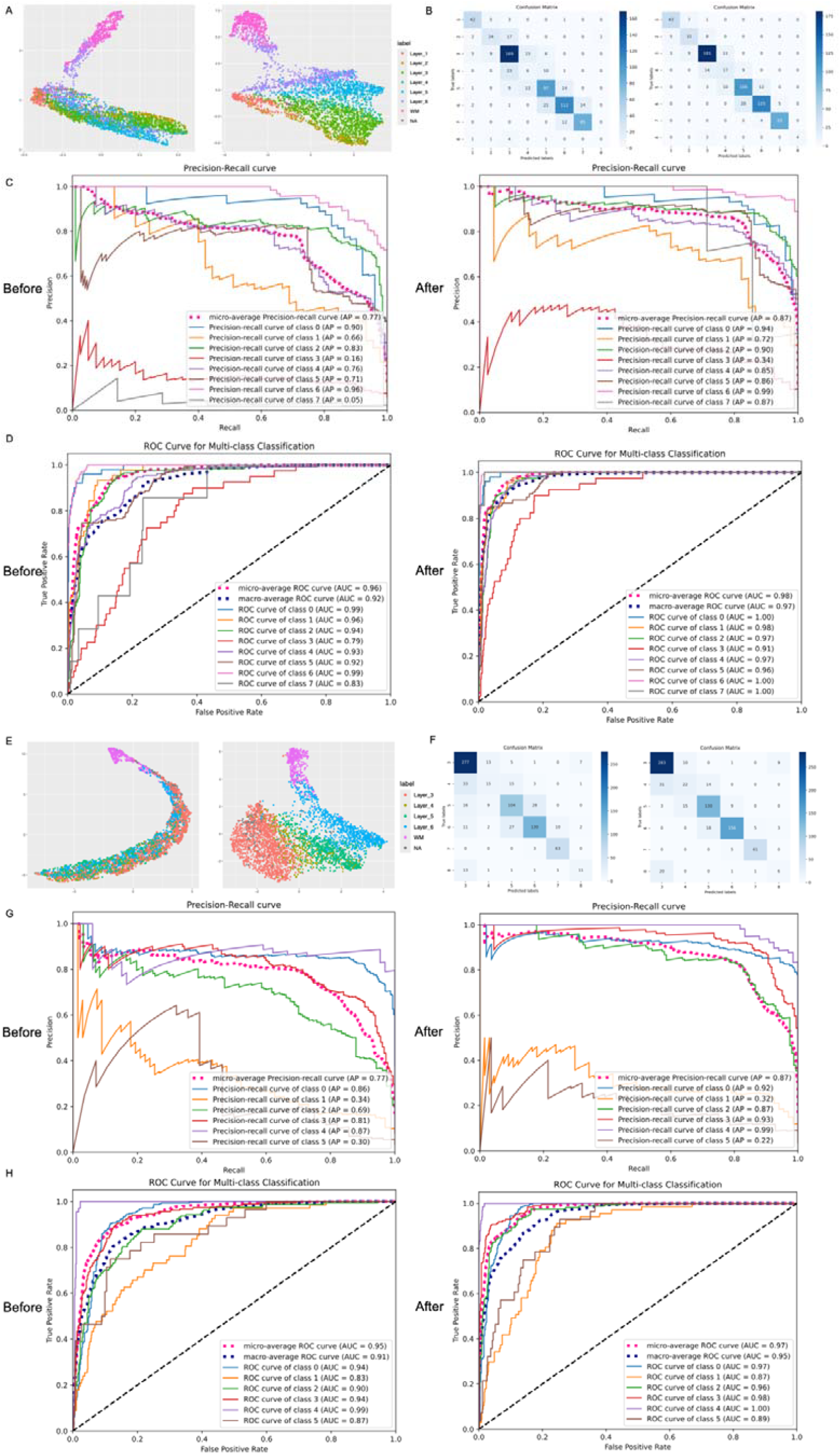
DiffusionST can enhance data quality. **(A)** A two-dimensional scatter plot is depicted illustrating the original top 4096 features alongside the reconstructed top 4096 features within the DiffusionST model for the 151673 slice. **(B)** Confusion matrix before and after imputation for the 151673 slice. **(C)** Precision-Recall curve of the 151673 slice. **(D)** ROC curve of the 151673 slice. **(E)** A two-dimensional scatter plot is presented, showcasing the original top 4096 features juxtaposed with the reconstructed top 4096 features within the DiffusionST model for the 151672 slice. **(F)** Confusion matrix before and after imputation for the 151672 slice. **(G)** Precision-Recall curve of the 151672 slice. **(H)** ROC curve of the 151672 slice.

### Comparison of the clustering accuracy of the imputation results of DiffusionST with other popular imputation algorithms and DiffusionST can adapt to high-resolution spatial transcriptomics

To further validate the applicability of DiffusionST for spatial transcriptomic data, we compared the data processed by the DiffusionST model with that repaired by five other popular imputation algorithms on twelve benchmark datasets. To ensure a fair comparison of the results after data imputation (repairing), we applied the same initialization parameters to the data imputed by the six tools and performed k-means clustering based on known cluster data. We compared the results using two-dimensional scatter plots and ARI accuracy of clustering labels with the true labels. We found that after imputing data from twelve datasets, the two-dimensional scatter plots generated using UMAP for the data processed by DiffusionST were more dispersed compared to the other five algorithms (as shown in **Figure 5A** and **Supplementary Figure S7-S12**). The two-dimensional scatter plots of other algorithms exhibited continuity but with lower clustering accuracy. In **Figure 5B**, we found that DiffusionST ranked first in clustering accuracy among the twelve datasets compared to the other five algorithms. We found that the clustering accuracy of DiffusionST after repairing data was higher than 0.2 in nine datasets. ALRA, MAGIC, and kNN_smoothing had clustering accuracy between 0.1 and 0.2 in twelve datasets after imputatio<u>n</u>, while DCA had clustering accuracy lower than 0.1 in two datasets after imputation, and SAVER had clustering accuracy lower than 0.1 in six datasets after imputation. We also used spectral clustering and the DBSCAN algorithm to test the clustering results after different algorithmic feature processing and found that spectral clustering yielded the highest accuracy on 12 datasets after DiffusionST feature processing. However, DBSCAN does not seem well-suited for clustering spatial transcriptomics data, as many clustering accuracies were zero (**see Supplementary Figure S13-S14**). In **Figure 5C**, DiffusionST can also be adapted to high-resolution data, with continuous clustering results.

**Figure 5.**
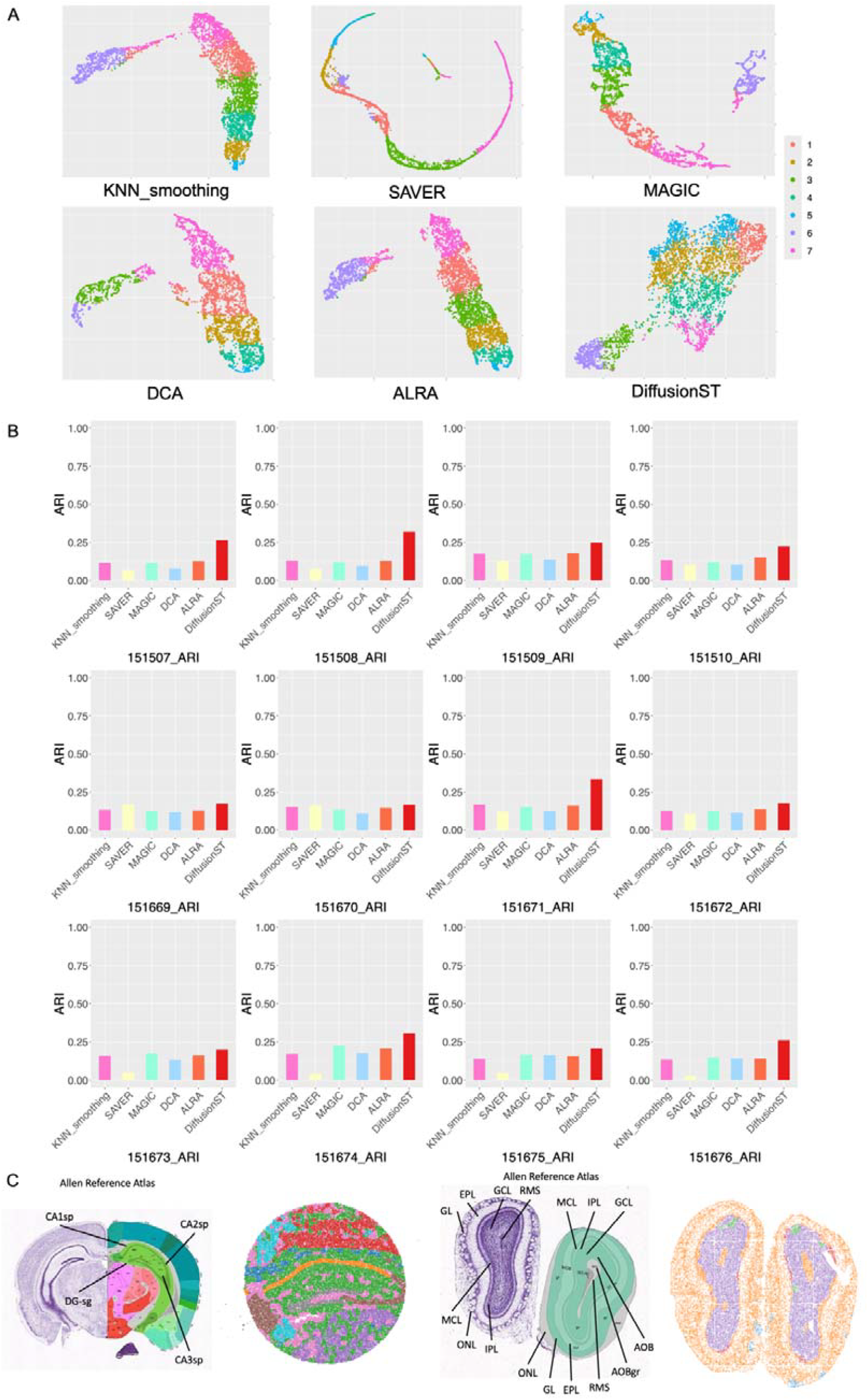
DiffusionST is compared with other mainstream imputation algorithms and can be adapted to high-resolution spatial transcriptomics. **(A)** A two-dimensional scatter plot of the 151673 slice data after imputation using the DiffusionST model compared to other popular imputation models. **(B)** Comparison of ARI scores between DiffusionST and five other imputation algorithms using the k-means algorithm. **(C)** DiffusionST shows continuity in clustering results for high-resolution data.

### DiffusionST provides a detailed dissection of spatial domains within cancer tissues

We applied the DiffusionST to the breast cancer spatial transcriptomic data to test its ability to dissect the spatial domain within the cancer tissues. In slice CID4465[34], we found that the clustering results obtained using DiffusionST were consistent with the labeling provided by the pathologist of the original dataset (as shown in **Figure 6A**). Additionally, although the performance of all algorithms was not ideal due to the data quality of this slice, DiffusionST achieved the highest ARI and provided a more detailed classification compared to STAGATE and GraphST (as shown in **Figure 6A**). In the region labeled as a tumor by the original annotation, we identified three domains that exhibited significant differences in terms of tumor progression and immune response.

**Figure 6.**
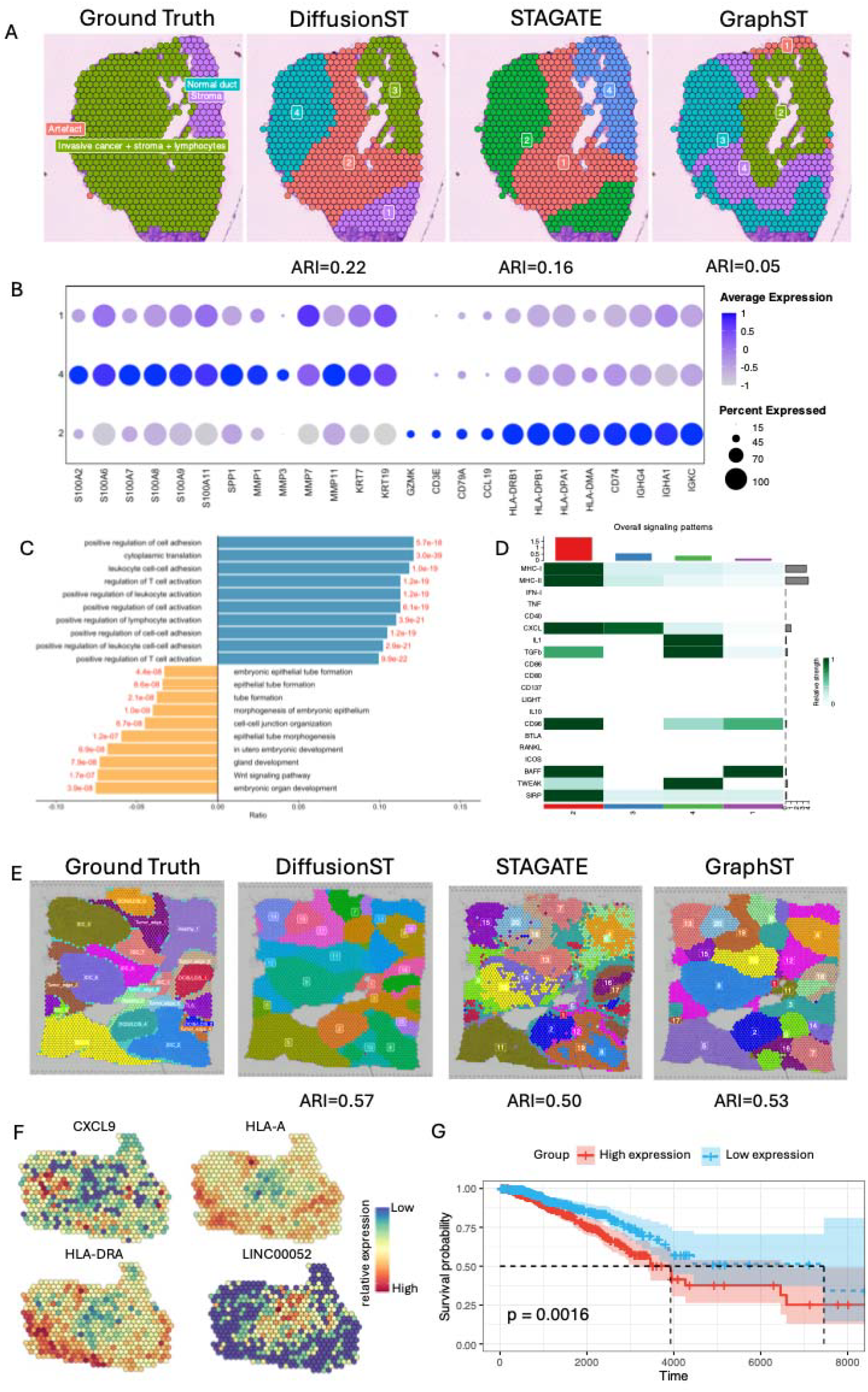
DiffusionST provides a detailed dissection of spatial domains. **(A)** Visium spatial transcriptomics data CID4465 labeled by pathologists, DiffusionST, STAGATE and GraphST. **(B)** Bubble map of marker genes expression within 3 tumor domains. **(C)** Biological pathways of GO enrichment analysis. Blue represent domain2 enriched GO terms and orange represent domain4 enriched GO terms. P-values are labeled at side of the bar. **(D)** Heatmap of immune-related cell-cell communction pathways of each domain. **(E)** Visium spatial transcriptomics data of breast cancer annotated by pathologists, DiffusionST, STAGATE and GraphST. **(F)** Visualization of marker gene expression of domain 14, 17, 19. **(G)** Kaplan-Meier plot showing better prognosis in patients with low average expression of domain 19 marker genes. The high expression group and low expression group are identified by the mean expression level of the prognostic marker genes from domain 19 (log rank test, p = 0.0016).

Bubble map showed the expression of marker genes in those 3 domains. We found that extracellular matrix reconstruction-related genes such as S100A2, S100A6 and MMP1 are high expressed in domain 4, and immune infiltration-related genes, including HLA-DRB1, HLA-DPB1 and IGHG4, are highly expressed in domain 2 (as shown in **Figure 6B**). These expression patterns demonstrate that in domain 4, extracellular matrix reconstruction occurs, suggesting a higher degree of tumor progression, and in domain 2 the immune response pathway is activated[35,36]. Domain 1 exhibits a moderate degree of tumor progression, as well as a weaker immune response. Next, we performed differential gene expression analysis between domain2 and domain4, and the differential expressed genes were subjected to Gene Ontology enrichment analysis (as shown in **Figure 6C**). We found that biological pathways enriched in domain 4 are related to epithelial cell developmental processes. In addition we performed cell cell communction analysis, which showed that the signal of immune-related pathways was significantly higher in domain4 than in other domains (as shown in **Figure 6D**).These results suggested that in domain4 the tumor cells exhibit a high activation, which may contribute to the tumor proliferation and metastasis[35-[37]. In contrast, pathways associated with immune response were activated in domain2, indicating a stronger immune response in this region.

Collectively, DiffusionST can provide more detailed insights into cancer spatial transcriptomic data, which can help us investigate tumor heterogeneity in greater detail and thus enhance cancer treatment strategies. We also conducted analysis and discussion on another breast cancer spatial transcriptomics dataset (**Supplementary Note S5**).

## Discussion

In this paper, we propose DiffusionST, which achieves high clustering accuracy on 12 benchmark low-resolution spatial transcriptomics datasets. The high clustering accuracy of the imputed data by DiffusionST can be attributed to five main aspects. Firstly, selecting appropriate feature genes and preprocessing the data. Secondly, DiffusionST uses the GCN model with ZINB loss function to learn the spatial location information and gene expression matrix of spatial transcriptomics for data denoising. Thirdly, the denoised data is imputed by using a U-net model with diffusion and multi-head attention mechanisms to learn the data distribution. Fourthly, compared to Generative Adversarial Networks (GANs), Diffusion Models gradually learn the increasing complexity of data distributions (from noise to data), producing more diverse and stable samples while avoiding the issue of mode collapse. The generation process is based on explicit probabilistic modeling, progressively approximating the data distribution from noise, offering strong theoretical interpretability. Finally, the data enhancement matrix is clustered using “mclust” package[38]. DenoiseST can effectively address the challenges posed by large datasets. It is noteworthy that testing on the 10x Visium dataset, which includes approximately 4,000 spots, requires 30 minutes of computation time on a server equipped with an AMD EPYC 7502P 32-core processor CPU and an NVIDIA GA100 GPU. Additionally, evaluation of a high-resolution dataset containing approximately 50,000 spots on the same server configuration takes 2 hours of computation time. DiffusionST is scalable and can reduce computation time by adjusting the iteration count parameter in the code.

In summary, DiffusionST achieved high accuracy in imputation on twelve benchmark datasets. DiffusionST can effectively handle dropout noise in spatial transcriptomics data. Compared to five other popular imputation algorithms, DiffusionST has better imputation performance. DiffusionST demonstrates superior resistance to random initialization and can be scaled to handle datasets with tens of thousands of spots. Additionally, DiffusionST can identify delicate regions in spatial transcriptomics data, such as those found in breast cancer tissue slices, which helps us to further understand tumor heterogeneity and is expected to contribute to the development of new therapeutic strategies for cancer in the future.

## Methods

### Method overview

DiffusionST first requires the selection of feature genes and data preprocessing for spatial transcriptome data. Spatial transcriptome data provide gene expression data and spatial location information. We reviewed recent relevant research literature, and the threshold for selecting feature genes ranges from the top 3000 to the top 5000. To facilitate the formation of a spot vector into a matrix, we used the “seurat_v3” function in the scanpy[30] package to filter and normalize the raw gene expression genes, with a threshold of 4096 for selecting feature genes. We can obtain the preprocessed gene expression counts matrix *M*.

Next, we utilized the nonlinear model of the DenoiseST[39] tool to perform feature learning on the preprocessed genes, employed a GCN model to integrate the features of each spot with those of its neighboring spots, and used a ZINB model for feature learning of each gene to better fit the ZINB distribution. Finally, obtain the embedded matrix *x*_0_.

### The data Imputation process of the Diffusion model

Using a diffusion model to further enhance the data quality of the input matrix *H* obtained from the ZINB embeddings integrated by GCN. Firstly, we convert each embedded spot vector into a 64 × 64 image format. Secondly, input each spot’s image sequentially into the Diffusion image restoration model. The specific reasons for selecting 4096 features are shown in Table S3. The diffusion model includes two processes: the forward process and the reverse process, with the forward process also known as the diffusion process. Both the forward process and the reverse process are parameterized Markov chains, where the reverse process can be used to generate data, and here we will use variational inference for modeling and solving.

The diffusion process refers to the gradual addition of Gaussian noise to the data until it becomes random noise. For the embedded data *x*_0_∼ *q*(x_0_), each step in the diffusion process, which comprises steps in total, involves adding Gaussian noise to the data *x*_*t−*1_ obtained from the previous step as follows:

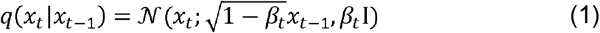

Here, 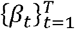 is the variance used at each step, ranging between 0 and 1. For the diffusion model, the variance settings for different steps are often referred to as the variance schedule or noise schedule. Typically, later steps use larger variances, which satisfies *β*_1_ < *β*_2_ <…< *β*_*T*_. Under a well-designed variance schedule, if the diffusion steps are sufficiently large, the result will completely lose the original data and become random noise. Each step of the diffusion process generates a noisy data spot, and the entire diffusion process forms a Markov chain:

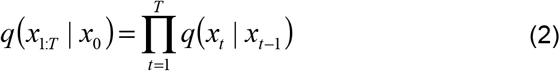

An important feature of the diffusion process is that we can directly sample *x*_0_ at any step based on the original data *x*_*t*_ : *x*_*t*_∼ q(x_*t*_|x_0_). Here, we define *a*_*t*_ = 1 − *β*_*t*_ and 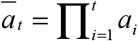, and through the reparameterization trick (similar to VAE), we have the following derivation process:

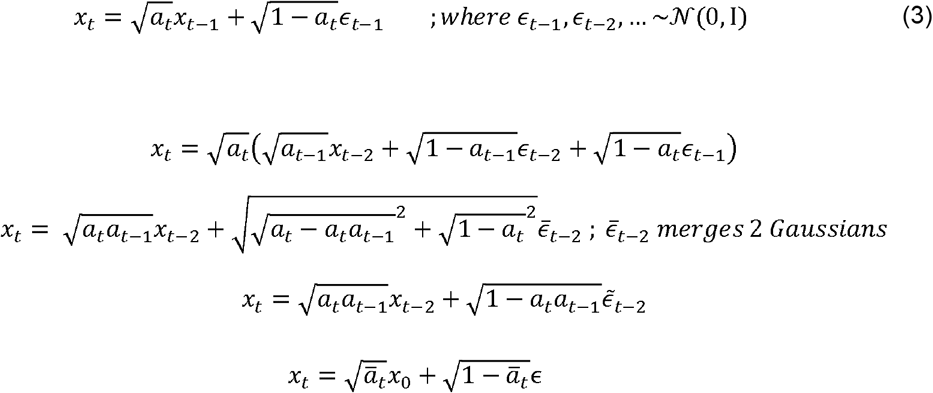

The above derivation utilized two Gaussian distributions with different variances 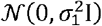 and 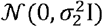, and their sum equals a new Gaussian distribution 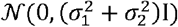. After reparameterization, we obtain:

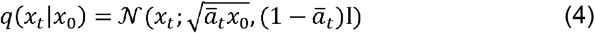

### Reverse process

The diffusion process adds noise to the image, while the reverse process is a denoising process. If we know the true distribution *q*(*x*_*t*-1_|*x*_*t*_) at each step of the reverse process, we can start from random noise *x*_*T*_∼ *𝒩* (0, 1) and gradually denoise to generate a real sample. Therefore, the reverse process is also a data generation process.

Estimating the distribution requires the entire training sample, and we can use neural networks to estimate these distributions. Here, we define the reverse process as a Markov chain[40,41] composed of a series of Gaussian distributions parameterized by neural networks:

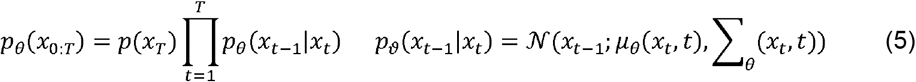

Among them, *p*(*x*_*T*_) = *𝒩* (*x*_*T*_; 0,?), and *p*_0_(*x*_*t*-1_|*x*_*t*_) is a parameterized Gaussian distribution. Their mean and variance are given by the trained network *µ*_0_ (*x*_*t*_, *t*) and ∑_*θ*_(*x*_*t*_, *t*).

Compared to VAE, the latent variables in diffusion models have the same dimensions as the original data, and the encoder (i.e., the diffusion process) is fixed. Since the diffusion model is a latent variable model, we can use variational inference to obtain the variational lower bound (VLB, also known as ELBO) as the maximization objective. Here we have:

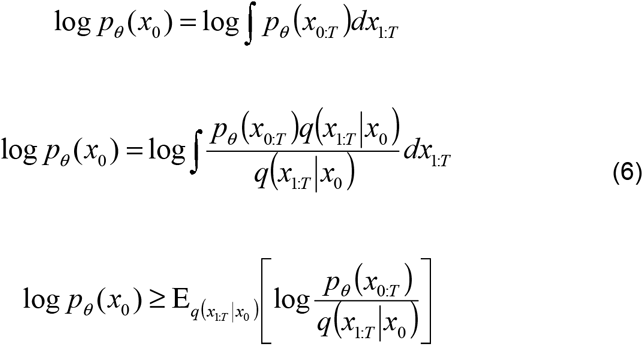

The final step utilizes Jensen’s inequality. For network training, the training objective is the negative VLB:

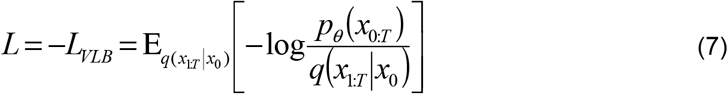

We further decompose the training objective to obtain:

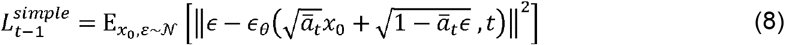

Since noise and original data are of the same dimensionality, we can choose to use an autoencoder architecture as the noise prediction model. The model we used is a U-Net model based on residual blocks and attention blocks. U-Net belongs to the encoder-decoder architecture, where the encoder is divided into different stages, each stage containing down-sampling modules to reduce the spatial size of features. Then, the decoder and encoder are opposite, gradually recovering the compressed features of the encoder. In the decoder module, U-Net also introduces skip connections, concatenating the same-dimensional features obtained from the encoder, which is beneficial for network optimization. The U-Net used by DDPM includes 2 residual blocks in each stage, and some stages also include self-attention modules to increase the network’s global modeling capability. Additionally, the diffusion model requires a noise prediction model. In practice, we can add a time embedding (like the position embedding in transformers) to encode the timestep into the network, so that we only need to train a shared U-Net model. Specifically, DDPM introduces time embeddings in each residual block. We used a U-net model architecture. We changed the attention layer, using multi-head attention[42], and chose to use four attention heads instead of one (while maintaining the same total number of channels). We calculate the similarity between the pseudo-images generated by the diffusion model for each spot and the original images. The two pseudo-images most like the processed spot are selected and averaged to replace the processed spot. Finally, clustering is performed on the enhanced data following data augmentation.

## Key Points

- DiffusionST effectively imputes data from twelve benchmark datasets, validated by improved clustering performance.
- It outperforms six popular spatial transcriptomic clustering algorithms and five scRNA-seq imputation methods in clustering accuracy.
- The model demonstrates robustness against noise, significantly enhancing data quality under random dropout noise.
- DiffusionST is well-suited for high-resolution spatial transcriptomics, with applications in survival analysis and cell-cell communication studies.
- These results highlight DiffusionST’s efficacy and value in processing noisy spatial transcriptomic data.

## Availability of Data and Materials

The source code package is freely available at https://github.com/cuiyaxuan/DiffusionST. The datasets used in this study can be found at https://drive.google.com/drive/folders/1qgn2UKpu4q14ysCoCKjWYVEHXIzHNoqq?usp=drive_link. The details of the data description are shown in **Supplementary Note S1**. Additionally, we have set up a web tutorial to make it easier for users to access (https://diffusionst-tutor1.readthedocs.io/en/latest/).

## Acknowledgement

Leyi Wei and Xiucai Ye conceived and supervised the project. Yaxuan Cui designed the DiffusionST model and implemented it. Yaxuan Cui and Yang Cui contributed to the writing of the manuscript. Yang Cui led the biological data analysis. Yaxuan Cui and Yang Cui assembled the figures for the thesis. Yaxuan Cui and Ruheng Wang revised the paper. Feifei Cui, Zilong Zhang, Yan Chen and Hua Shi reviewed and checked the paper. Kenta Nakai and Tetsuya Sakurai provided the experimental equipment and computational resources.

